# Regulation of beta-amyloid production in neurons by astrocyte-derived cholesterol

**DOI:** 10.1101/2020.06.18.159632

**Authors:** Hao Wang, Joshua A. Kulas, Heather A. Ferris, Scott B. Hansen

## Abstract

Alzheimer’s Disease (AD) is characterized by the presence of β-Amyloid (Aβ) plaques, tau tangles, inflammation, and loss of cognitive function. Genetic variation in a cholesterol transport protein, apolipoprotein E (apoE), is the most common genetic marker for sporadic AD. *In vitro* evidence suggests apoE links to Aβ production through nanoscale lipid compartments (also called lipid rafts), but its regulation *in vivo* is unclear. Here we use super-resolution imaging in mouse brain to show apoE utilizes astrocyte-derived cholesterol to specifically traffic neuronal amyloid precursor protein (APP) into lipid rafts where it interacts with β- and γ-secretases to generate Aβ-peptide. We find that targeted deletion of astrocyte cholesterol synthesis robustly reduces amyloid and tau burden in a mouse model of AD. Treatment with cholesterol-free apoE or knockdown of cholesterol synthesis in astrocytes decreases cholesterol levels in cultured neurons and causes APP to traffic out of lipid rafts where it interacts with α-secretase and gives rise to soluble APPα (sAPPα), a neuronal protective product of APP. Changes in cellular cholesterol have no effect on α-, β-, and γ-secretase trafficking, suggesting the ratio of Aβ to sAPPα is regulated by the trafficking of the substrate, not the enzymes. Treatment of astrocytes with inflammatory cytokines IL-1β, IL-6 and TNF-α upregulates the synthesis of cholesterol in the astrocytes. We conclude that cholesterol is kept low in neurons to inhibit Aβ formation and enable astrocyte regulation of Aβ formation by cholesterol regulation.

**Highlights:** ApoE regulates amyloid precursor protein localization to rafts and its exposure to α-vs. β-secretase.

α-, β-, and γ-Secretases are activated by substrate presentation.

ApoE specifically transports astrocyte cholesterol to neurons.

Astrocyte cholesterol synthesis disruption prevents Alzheimer’s-associated amyloid pathology in mice.

## INTRODUCTION

Alzheimer’s disease (AD), the most prevalent neurodegenerative disorder, is characterized by progressive loss of cognitive function and the accumulation of amyloid β (Aβ) peptide and phosphorylated tau^1^. Amyloid plaques are composed of aggregates of Aβ peptide, a small hydrophobic protein excised from the transmembrane domain of amyloid precursor protein (APP) by proteases known as beta-(β-) and gamma-(γ-) secretases (Figure S1a). In high concentrations, Aβ peptide forms toxic species and aggregates to form Aβ plaques^2,3,4^. The non-amyloidogenic pathway involves a third enzyme, alpha-(α-) secretase, which generates a soluble APP fragment (sAPPα), helps set neuronal excitability in healthy individuals^5^ and does not contribute to the generation of amyloid plaques. Therefore, by preventing Aβ production, α-secretase-mediated APP cleavage is protective. Strikingly, both pathways are finely regulated by cholesterol^6^(Figure S1b).

**Figure 1.**
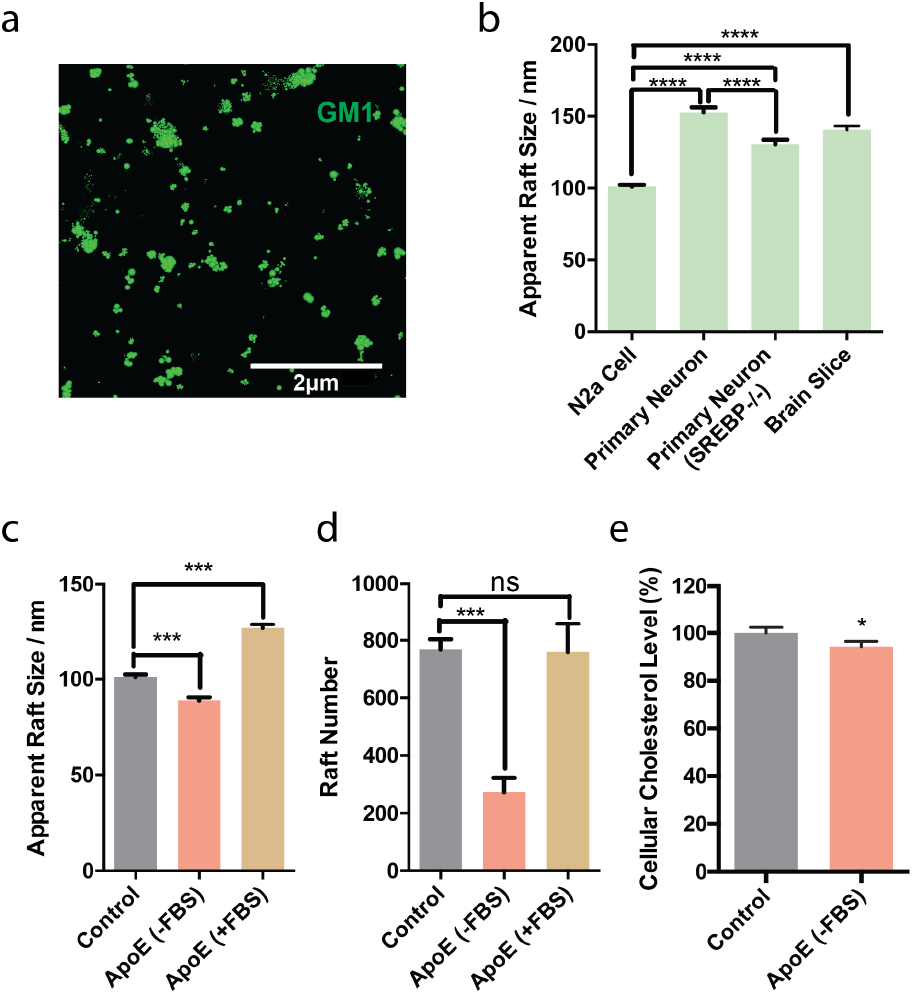
Depletion of astrocyte cholesterol decreases neuronal lipid raft size. (a) dSTORM super resolution imaging on brain slices showing lipid raft nano-structure in cell membrane. Scale bar is 2 μm. (b) Comparison of apparent raft length in neuroblastoma 2a (N2a) cells, primary neurons cultured with astrocytes with and without (SREBP-/-) cholesterol synthesis and brain slices. (c) Quantitation of raft lengths indicates changes in raft structure after apoE treatment. (d) Quantitation of GM1 clusters shows apoE decreased the number of rafts under low cholesterol conditions, while apoE treatment with high environmental cholesterol doesn’t affect raft number. Data are expressed as mean ± s.e.m., one-way ANOVA. (e) Exposure of N2a cells to apoE removes cholesterol from cellular membranes as measured by a fluorescent based cholesterol assay. Data are expressed as mean ±s.e.m., two-sided Student’s t-test; *P<0.05, ***P<0.001, ****P<0.0001.

In cellular membranes, cholesterol regulates the formation of lipid domains (also known as lipid rafts) and the affinity of proteins to lipid rafts^7^ including β-secretase and γ-secretase^8–10^. α-Secretase does not reside in lipid rafts, rather α-secretase is thought to reside in a region made up of disordered polyunsaturated lipids^11^. The location of APP is less clear. In detergent resistant membranes (DRMs) studies, it primarily associates with lipid from the disordered region, although not exclusively^8,10,12,13^. Endocytosis is thought to bring APP in proximity to β-secretase and γ-secretase and this correlates with Aβ production. Cross linking of APP with β-secretase on the plasma membrane also increases Aβ production, leading to a hypothesis that raft movement in the membrane contributes to APP processing^11,14^ (Figure S1a). Testing this hypothesis *in vivo* has been hampered by the small size of lipid rafts (often <100 nm), which is below the resolution of light microscopy.

Super resolution imaging has emerged as a complimentary technique to DRM’s, with the potential to interrogate raft affinity more directly in a native cellular environment^15^. We recently employed super resolution imaging to establish a membrane mediated mechanism of general anesthesia^16^. In that mechanism, cholesterol causes lipid rafts to sequester an enzyme away from its substrate. Removal of cholesterol then releases and activates the enzyme by giving it access to its substrate (Figure S1c)^7,17^. A similar mechanism has been proposed to regulate the exposure of APP to its cutting enzymes^11,14,18–20^.

Neurons are believed to be the major source of Aβ in normal and AD brains^21,22^. In the adult brain, the ability of neurons to produce cholesterol is impaired^23^. Instead, astrocytes make cholesterol and transport it to neurons with apolipoprotein E (apoE)^24–26^. Interestingly, apoE, specifically the e4 subtype (apoE4), is the strongest genetic risk factor associated with sporadic AD^27,28^. This led to the theory that astrocytes may be controlling Aβ formation through regulation of lipid raft function^11,14,18^, but this has not yet been shown in the brain of an animal. Here we show that astrocyte-derived cholesterol controls Aβ formation *in vivo* and links inflammatory cytokines, apoE, Aβ, and plaque formation to a single molecular pathway.

## RESULTS

### Characterization of astrocyte derived cholesterol on neuronal raft formation

To establish a role for Aβ regulation by astrocytes *in vivo*, we first labeled and imaged monosialotetrahexosylganglioside1 (GM1) lipids in wild type mouse brain slices. GM1 lipids reside in cholesterol dependent lipid rafts and bind cholera toxin B (CTxB) with high affinity^29,30^. These GM1 domains are separate from phosphatidylinositol 4,5 bisphosphate (PIP2) domains, which are polyunsaturated and cholesterol independent^17,31,32^. We labeled GM1 domains (i.e. lipid rafts) from cortical slices with Alexa Fluor 647 conjugated fluorescent CTxB, and imaged with confocal and super-resolution direct stochastical optical reconstruction microscopy (dSTORM)^33–36^. dSTORM is capable of visualizing nanoscale arrangements (i.e. sub-100 nm diameter lipid domain structures^37^) in intact cellular membranes.

CTxB appeared to label most cell types in cortical brain slices (Figure S2a, green shading). In neurons, labeled with a neuron specific antibody against neurofilament medium chain (NFM) protein, CTxB can be seen outlining the plasma membrane (outside of the cell) as opposed to NFM which labels throughout the cells (Figure S2a right panels).

Next, we investigated the role of cholesterol on the relative size and number of neuronal GM1 domains in brain tissue (Figure 1a) using dSTORM (a ~10-fold increase in resolution compared to confocal). In wild type mouse cortical tissue GM1 domains averaged ~141 nm in apparent diameter, slightly smaller than the apparent size in primary neurons (~150 nm diameter) (Figure 1b). Cultured neuroblastoma 2a (N2a) cells exhibited the smallest rafts by far, on average only 100 nm in apparent diameter. All of the mammalian cells had domains larger than intact fly brain which had an apparent diameter of ~90 nm^16^. CTxB is pentadentate and can affect the absolute size, here we used the numbers merely as a relative comparison of size under identical conditions (see supplemental discussion).

Next we compared the size of neurons cocultured with cholesterol deficient astrocytes. When cholesterol was depleted from astrocytes, the raft size of primary neurons was significantly reduced (130 nm). We depleted astrocyte cholesterol by SREBP2 gene ablation (SREBP2-/-). SREBP2 is an essential regulator of cholesterol synthesis enzymes^38^ and was specifically knocked out in astrocytes using an ALDH1L1 promoter-driven Cre recombinase^39^. The observed effect of astrocyte SREBP2 ablation on neurons suggests decrsed cholesterol transport to neurons—presumably through apoE.

To confirm that astrocytes could regulate neuronal GM1 domain formation through apoE, we added purified apoE to cultured primary neurons and N2a cells. The apoE (human subtype 3) derived from *E. coli* was devoid of cholesterol, as prokaryotes do not make cholesterol. To provide a source of cholesterol to the apoE, we added 10% fetal bovine serum (FBS), a common source of mammalian lipids, including ~310 μg/mL of cholesterol containing lipoproteins. ApoE can both load and unload cholesterol from cells, including neurons^40–42^. Importantly, apoE is not present in FBS^43^. Cells were treated acutely (1 hr) with 4 μg/mL of apoE, a physiologically relevant concentration seen in cerebral spinal fluid^44^.

Loading cells with cholesterol (apoE +FBS) caused an increase in the apparent raft diameter from 100 nm to 130 nm in N2a cells (Figure 1c). When cholesterol was unloaded, i.e., effluxed (ApoE - FBS), the apparent size and number of GM1 domains decreased (Figure 1c-d). Binning rafts by large > 500 nm and small < 150 nm showed a clear shift toward smaller domains with apoE in low cholesterol and a clear shift to large domains with apoE in high cholesterol (Figure S2c-d). To confirm this result, we compared apoE treatment to treatment with methyl β-cyclodextrin (MβCD), a non-native chemical binder that extracts cholesterol from the plasma membrane and disrupts raft function^45^. MβCD caused a similar decrease as seen for cells with cholesterol effluxed (Figure S2f-g).

We also investigated apoE’s effect in modulating membrane cholesterol level and lipid raft integrity using an enzymatic raft disruption assay. The disruption assay uses phospholipase D (PLD)-as a reporter of raft disruption^17^. After 1 h incubation with apoE (-FBS) PLD activity significantly increased, confirming raft disruption (Figure S2e). We confirmed apoE decreased N2a total cellular cholesterol (Figure 1e) using a fluorescent cholesterol assay (see methods). These results suggest a functional role of apoE in extracting cellular cholesterol and disrupting lipid rafts in cellular membranes.

### Super resolution imaging of amyloid processing proteins in lipid rafts

Next, we sought to characterize a cellular function of cholesterol loading, i.e. the ability of GM1 domains to move APP towards or away from its hydrolyzing secretases. To establish the movement of APP to each of its processing enzymes, a, β, and γ,-secretases, directly in cellular membranes, we imaged N2a cells with dSTORM. APP was previously found in both raft and non-raft like fractions of DRMs^8,10,12,13^. Detergent resistant membranes are similar to GM1 lipid rafts. However, imaging APP is important since the amount of raft-like association in DRM’s could easily be affected by the detergent concentration used for preparation. We labeled GM1 domains with fluorescent CTxB and the amyloid proteins (APP, a, β, and γ,-secretases) with Cy3b labeled fluorescent antibodies and determined raft localization by pair correlation using DBSCAN of two color dSTORM images (Figure 2a).

**Figure 2.**
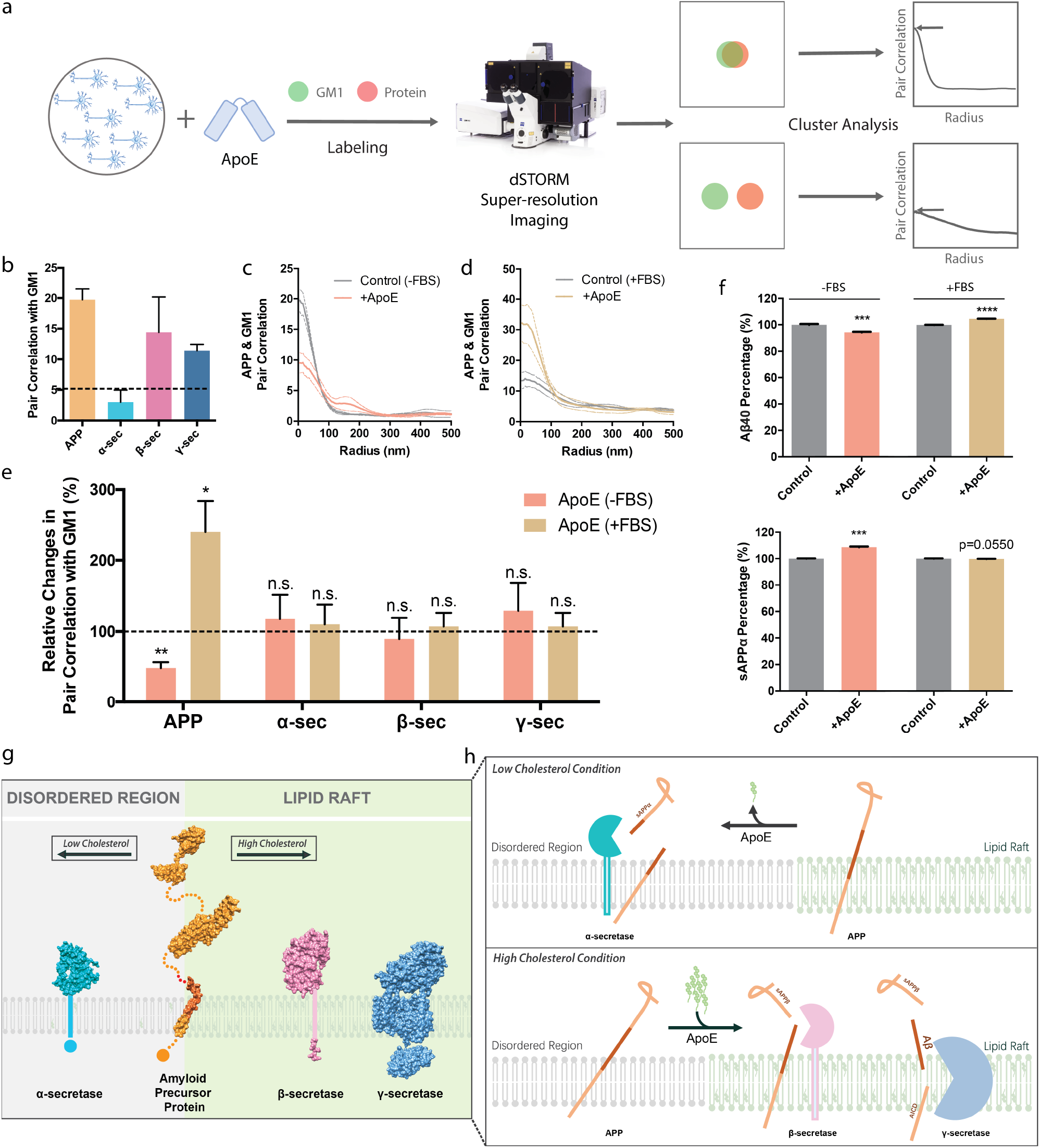
ApoE mediates Aβ production through cholesterol signaling. (a) Workflow for dSTORM superresolution. Cultured N2a cells (blue) are depicted in a dish. Cells were exposed to apoE with and without serum supplementation and fixed. GM1 lipids and amyloid proteins were fluorescently labeled (CTxB and antibody respectively) and imaged with super resolution (dSTORM). The proximity of the two labels (shown as red and green circles) was then determined by cluster analysis. Idealized pair correlations are shown for objects that strongly colocalize (top) and weakly co-localize (bottom) at a given radius. Pair correlations are unitless with 1 being little to no correlation and >5 a value typically significant in our experimental conditions. (b) α-secretase sits in the disordered region (low correlation with lipid raft) while APP, β- and γ-secretases are raft-associated. (c,d) Pair correlation analysis showing APP moving in (c) and out (b) of the GM1 rafts under high and low cholesterol conditions, respectively. (e) Under low cholesterol conditions (-FBS), APP co-localization with raft domains decreases markedly after apoE treatment. Under high cholesterol conditions (+FBS), apoE-mediated APP co-localization with raft domains increases (i.e., apoE induces APP to translocate into lipid rafts). α-, β- and γ-secretase localization do not respond to apoE-mediated raft modulation. (f) ELISA assay showing a shift from Aβ to sAPPα production after apoE treatment under low cholesterol condition and increased Aβ production under high cholesterol condition. Data are expressed as mean ± s.e.m. *P<0.05, **P<0.01, ***P<0.001, ****P<0.0001, two-sided Student’s t-test. (g, h) Because of raft disruption mediated by apoE, APP is ejected from lipid rafts into disordered regions under low cholesterol conditions, exposing it to α-secretase to be cleaved into non-amyloidogenic sAPPα. With high environmental cholesterol, apoE induced raft stabilization mediates more APP to be translocated into lipid rafts and cleaved by β- and γ-secretase, producing amyloidogenic Aβ peptides.

Figure 2b shows APP, β-secretase, and γ-secretase are associated with GM1 raft lipids (i.e. they exhibit strong pair correlation with CTxB labeled GM1 lipids at short distances (5nm)), consistent with experiments in DRMs. In contrast, a-secretase is not associated with GM1 domains (i.e. lacks pair correlation with CTxB) (Figure 2b). The pair correlation (a unitless number) ranged from 10-15 for APP, β-secretase, and γ-secretase while the pair correlation for a-secretase was <3, a value typical for little or no pair correlation, a result also consistent with previous experiments in DRMs^14^.

### Cellular regulation of APP exposure to α- and β-secretase by apoE

We then investigated the ability of apoE to mediate delivery of cholesterol to cell membranes and regulate Aβ production in N2a cells. We applied 4 μg/mL of human apoE isoform 3 (apoE3) purified from *E. coli* with and without FBS, and measured APP localization to lipid rafts with dSTORM. We found that when ApoE loaded cells with cholesterol (apoE +FBS), APP association with GM1 domains was almost two-fold higher than when apoE effluxed cholesterol (-FBS) (Figure 2c-e). In a control experiment, FBS alone (no apoE added), did not increase APP trafficking to rafts (Figure 2cd), confirming apoE is a specific and necessary mediator of cholesterol transport to the neuron. Moreover, apoE alone (under low cholesterol condition (-FBS)) decreased APP raft affinity (Figure 2b-c), confirming apoE can unload cholesterol in neuronal membranes. ApoE efflux was similar to MβCD treatment (Figure S2h-j)^11,14^.

Surprisingly, neither cholesterol loading nor unloading of N2a cells had a significant effect on α-, β-, or γ-secretase trafficking in or out of GM1 domains (Figure 2e). GM1 domains were analyzed pairwise for association with either α-, β-, or γ-secretase in an identical manner to APP. In contrast to APP, cholesterol unloading with apoE treatment was insufficient to move β-, or γ-secretase out of GM1 domains. Likewise, more cholesterol failed to increase the association of any of the secretases. α-Secretase, located in the disordered region, remained in the disordered region (Figures 2b,e), while β- and γ-secretases, which were already in the lipid rafts, did not increase their association with GM1 lipids in cholesterol loaded cells (Figures 2b,e). This suggests a mobile substrate (APP) moves between relatively static enzymes (secretases) in the plasma membrane. ELISA assay showed a shift from Aβ40 to sAPPα production under low cholesterol condition and high membrane cholesterol mediates increased Aβ40 generation (Figure 2f). Figures 2g-h summarize the substrate moving between the different secretases based on raft-associated apoE-dependent trafficking and the expected cleavage products based on localization. With low cholesterol, apoE moves APP to α-secretase generating more sAPPα (top panel). During high cholesterol apoE moves APP to β- and γ-secretase which produce neurotoxic Aβ peptide (bottom panel).

### The role of astrocyte cholesterol in regulating APP processing in cultured primary neurons

Since astocytes regulate cholesterol levels and raft formation in neurons (Figure 1), we hypothesized that astrocytes could directly control Aβ production in neuronal membrane. To test this hypothesis, we cultured primary cortical cells from mouse embryos (E17). In a first culture, we isolated just the neurons by sorting the cells with live cell fluorescence activated cell sorting (FACS) with Thy1.2, a cell surface marker of mature neurons. In a second culture we selected for neurons, but did not remove contaminating astrocytes resulting in a mixed culture. All cultures were treated with or without apoE, fixed, and labeled with appropriate fluorophores—the neurons were then imaged with dSTORM. We confirmed the presence of astrocytes in our mixed culture by staining for glial fibrillary acidic protein (GFAP). About 1% of the mixed cell population was labelled by the GFAP antibody (Figure S3f).

We found that incubation of purified cortical neurons (Thy1.2^+^, no astrocytes) with cholesterol-free human apoE dramatically decreased APP association with GM1 domains (>2-fold) (Figure 3b) in primary neurons. However, when we mixed astrocytes with neurons, the same apoE had the exact opposite effect. GM1 domains sequestered APP, evident by a 2.5-fold increase in APP association with GM1 lipids (Figure 3c). This suggests apoE delivers cholesterol from the astrocytes to the neurons and increases GM1 domain affinity for APP. Similar imaging of the secretases showed they remained immobile with α-secretase in the disordered region and β-secretase in GM1 domains (See Figure S3). Thus, astrocyte derived cholesterol is a potent signal that controls APP association with α- or β-secretase in the neuron, based on GM1 domain affinity.

**Figure 3.**
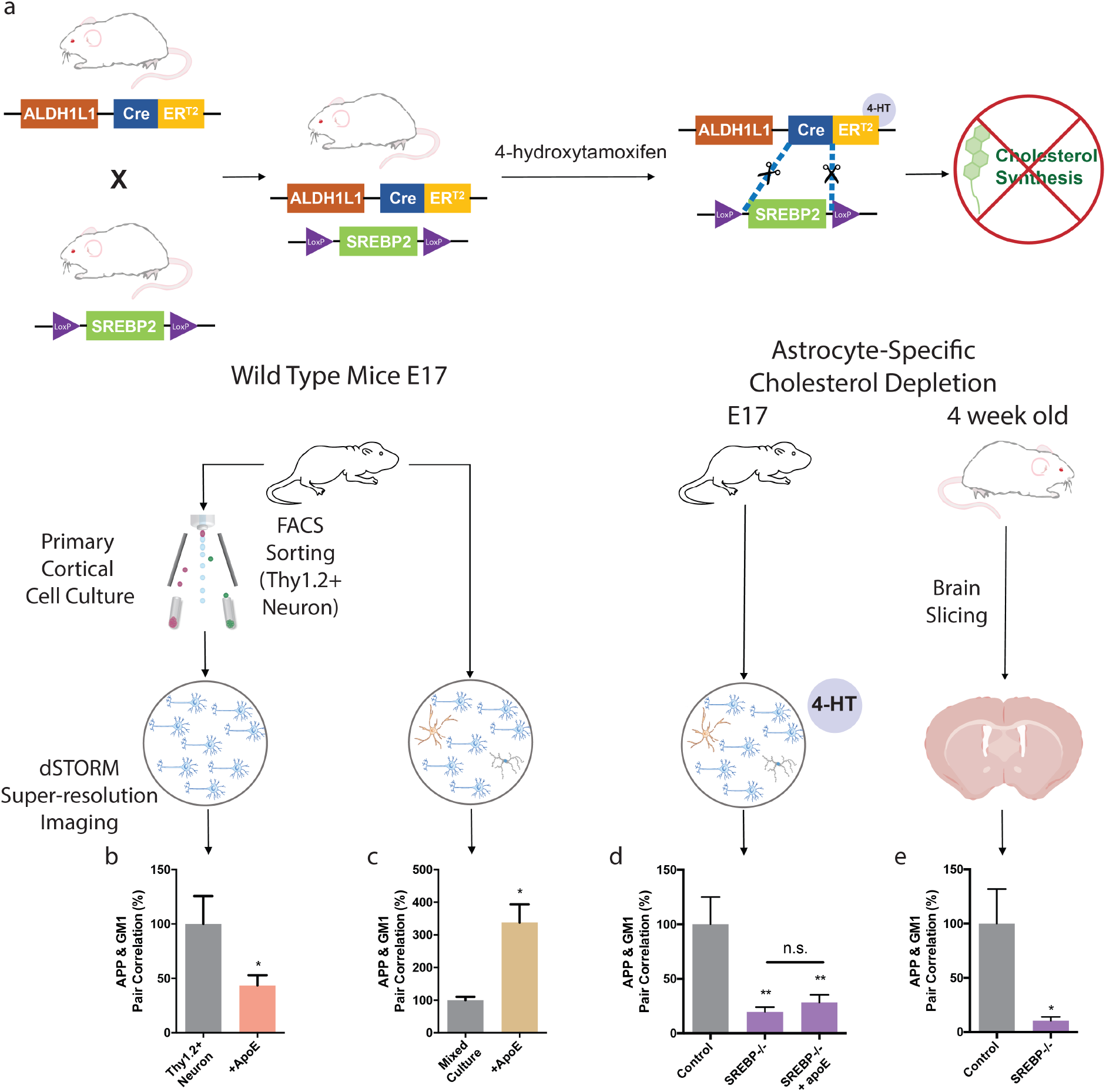
Astrocyte cholesterol regulates Aβ production in neurons. (a) Strategy for conditional knockout of SREBP2 in astrocytes using Cre-Lox recombination system. SREBP2 flox mice were crossed to ALDH1L1-specific Cre transgenic mice, which express Cre recombinase specifically in astrocytes when presented 4-hydroxytamoxifen (TMX). Cre promotes SREBP2 knockout and blocks cholesterol synthesis in astrocytes. (b-d) Cortical cells were isolated from embryonic day 17 mice and cultured for dSTORM imaging. (b) ApoE translocates APP from lipid rafts into disordered regions in pure neuronal population sorted with cell surface neuronal marker Thy1.2. (c) ApoE increases APP’s raft localization in primary neurons cultured in mixed population with glia cells including astrocytes. (d) Astrocyte-specific cholesterol depletion completely disrupts APP raft localization in neurons in cultured cells. (e) Astrocyte cholesterol depletion disrupts APP raft localization *in vivo* as demonstrated in SREBP2^fl/fl^GFAP-Cre^+/-^ mouse brain slices. Data are expressed as mean ± s.e.m., n=3-10, *P<0.05, **P<0.01, one-way ANOVA (d) or twosided Student’s t-test (b,c,e).

To further establish astrocyte derived cholesterol as the regulator of APP association with secretases, we specifically knocked down cholesterol synthesis in astrocytes by knocking out the SREBP2 gene using a tamoxifen-inducible Cre recombinase system (Figure 3a). SREBP2 is the master transcriptional regulator of the majority of enzymes involved in cholesterol synthesis. Mice with SREBP2 knocked out of astrocytes have reduced production of cholesterol in the astrocytes^46^ (Figure S4a). We cultured cortical neurons and astrocytes from E17 pups and treated them with 100 nM 4-hydroxytamoxifen for 2 days to induce knockout of astrocyte SREBP2. Three days after 4-hydroxytamoxifen removal the cells were fixed and labeled, and APP localization was imaged with dSTORM, identical to the FACS sorted neurons above (see diagrams Figure 3).

Figure 3d shows a highly significant decrease in APP pair correlation with GM1 domains in mixed cultures containing SREBP2 (-/-) astrocytes, similar to purified Thy1.2 positive neuronal cultures (Figure 3b). Control cells from Thy1.2^+^ neurons and SREBP2 (-/-) neuron/astrocyte cultures were very similar, the amplitude of pair correlation was ~10 for both cultures. 4-hydroxytamoxifen treatment did not significantly alter APP GM1 pair correlation in the absence of Cre recombinase (Figure S4b).

We next examined if astrocytes contribute cholesterol to promote APP presence in neuronal lipid rafts *in vivo*. Astrocyte cholesterol synthesis was disrupted in mice by crossing SREBP2 floxed mice to GFAP-Cre mice resulting in a homozygous deletion of SREBP2 from astrocytes^46^. At four weeks of age, brains were collected and APP-GM1 pair correlation was determined in neurons by dSTORM imaging. SREBP2 deletion in astrocytes resulted in a robust and significant reduction of APP-GM1 pair correlation compared to littermate controls (Figure 4a). This suggests that astrocyte cholesterol synthesis plays an essential role in maintaining cholesterol content in neuronal lipid rafts. These data further confirmed that astrocyte-derived cholesterol regulates APP translocation and association with secretases in the plasma membrane of neurons. Neurons appear incapable of increasing cholesterol synthesis to compensate for astrocyte deficiency in any appreciable way^47^.

**Figure 4.**
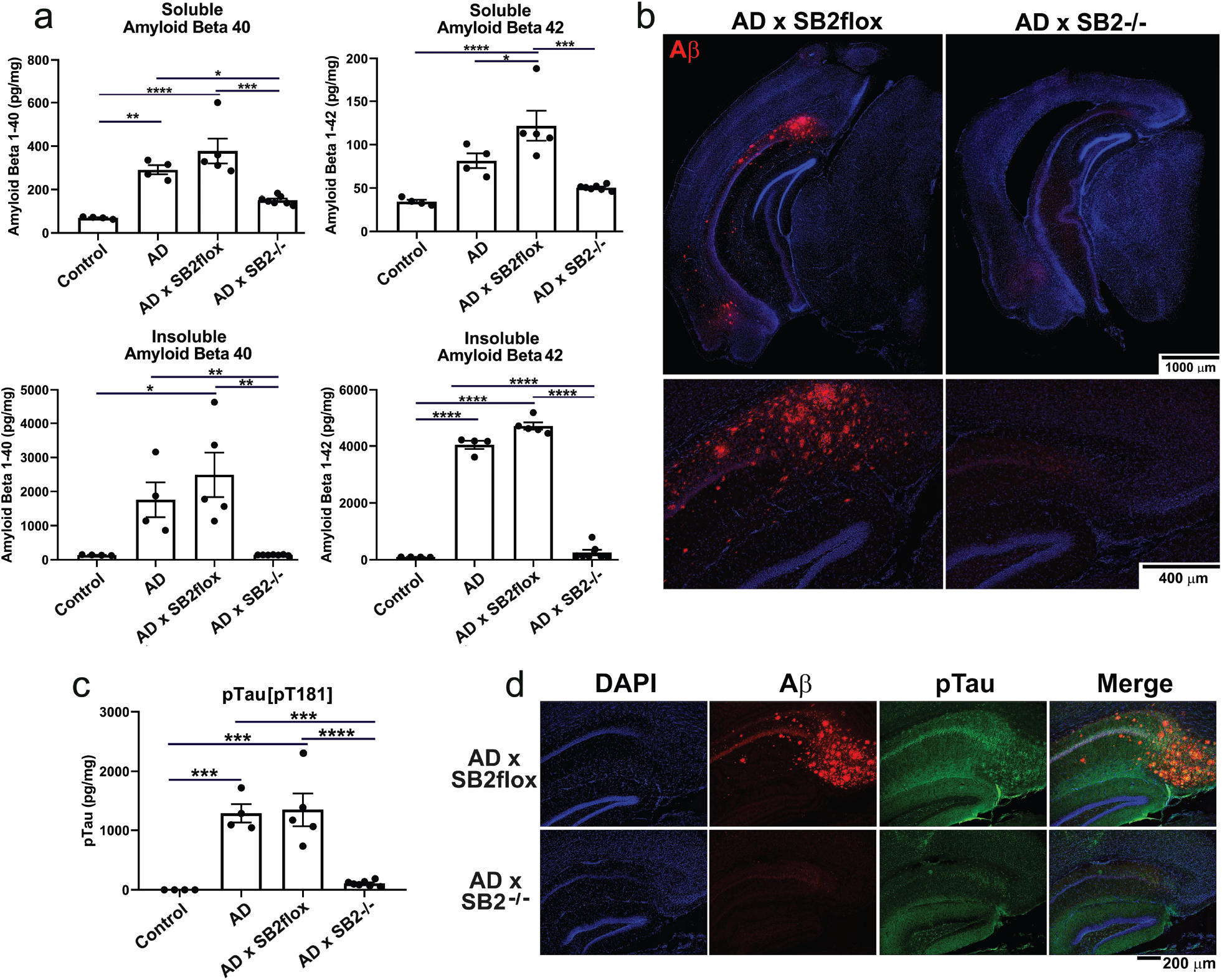
Loss of cholesterol synthesis in astrocytes blocks Aβ plaque formation and Tau phosphorylation *in vivo*. 3xTg-AD mice (AD) were crossed to SREBP2^fl/fl^GFAP-Cre^+/-^ mice (AD x SB2^-/-^) and aged to 60-weeks. (a) Hippocampus were isolated and soluble (RIPA-extracted) and insoluble (guanidine-extracted) amyloid-beta 40 and 42 species were measured by ELISA. (b) 60-week-old brain slices stained for human amyloid beta (red) and dapi (blue) in AD x SB2flox mice (left) and AD x SB2-/- mice (right). (c) pTau (phosphorylated threonine 181) levels were measured in 60-week-old hippocampus lysate by ELISA. (d) 60-week-old brain slices were immunostained for pTau (green) and amyloid beta (red) and the subiculum region of the hippocampus was imaged. Data are expressed as mean ± s.e.m, n=4-7, one-way ANOVA with Tukey’s post hoc analysis *P<0.05, **P<0.01, ***P<0.001, ****P<0.0001

### The *in vivo* role of astrocyte cholesterol on Aβ and plaque formation in mouse brain

As mentioned above, cholesterol is high in AD brains. Our model predicts attenuation of cholesterol in astrocytes should reduce the concentration of Aβ peptide formed *in vivo*. To test this hypothesis and to investigate the effect of astrocyte cholesterol on Aβ plaque formation in the intact brain we crossed 3xTg-AD mice, a well-established mouse model of AD^48^, with our SREBP2^flox/flox^ GFAP-Cre mice^46^ to generate an AD mouse model lacking astrocyte cholesterol. *In vivo*, Aβ exists in several forms, with Aβ1-40 being the most abundant and Aβ1-42 most closely associated with Alzheimer’s disease pathology. The 3xTg-AD mouse expresses transgenes for mutant human APP and mutant presenilin 1 enzyme, resulting in a significant increase in Aβ40 and Aβ42 production.

Using ELISAs, we measured human APP transgene derived Aβ40 and Aβ42, in both the RIPA soluble and insoluble fractions in the hippocampus of aged control and transgenic AD mice. Hippocampus tissue from wild type control mice was assessed to determine the nonspecific background of the Aβ signal in each ELISA assay. Knockout of SREBP2 in astrocytes of 3xTg-AD mice (AD x SB2-/-) reduced soluble Aβ40 and Aβ42 levels in hippocampus of 60-week-old mice by 2-fold, to levels only slightly higher than wildtype controls (Figure 4a) suggesting a near complete loss of amyloidogenic processing of APP. More impressively, insoluble Aβ40 and Aβ42 were almost entirely eliminated from AD x SB2-/- hippocampus (Figure 4a). This was verified by immunofluorescence staining demonstrating an absence of amyloid plaques in AD x SB2-/- mice (Figure 4b). Of note, total APP transgene expression was not impacted by loss of astrocyte SREBP2 (Figure S6). To further confirm that the reduced Aβ plaque burden in our model was due to reduced amyloidogenic processing, we performed mixed primary cultures of astrocytes and neurons from these 3xTg crosses. In agreement with our *in vivo* observation, we found that astrocyte SREBP2 deletion modestly increased sAPPα production and robustly reduced sAPPβ generation without changing total APP abundance (Figure 5a-c). The 3xTg-AD model also expresses a transgene for human tau protein, another key component of Alzheimer’s pathology. Tau is hyperphosphorylated in Alzheimer’s disease and this is thought to be downstream of amyloidbeta accumulation^49^. It has also been proposed that cholesterol directly regulates tau phosphorylation ^50,51^. In agreement with both of these hypotheses, phosphorylation of tau at the key T181 residue, but not total tau, is eliminated *in vivo* in the AD x SB2-/- hippocampus and reduced in mixed cultures *in vitro* (Figure 4c,d, Figure 5c, Figure S6).

**Figure 5.**
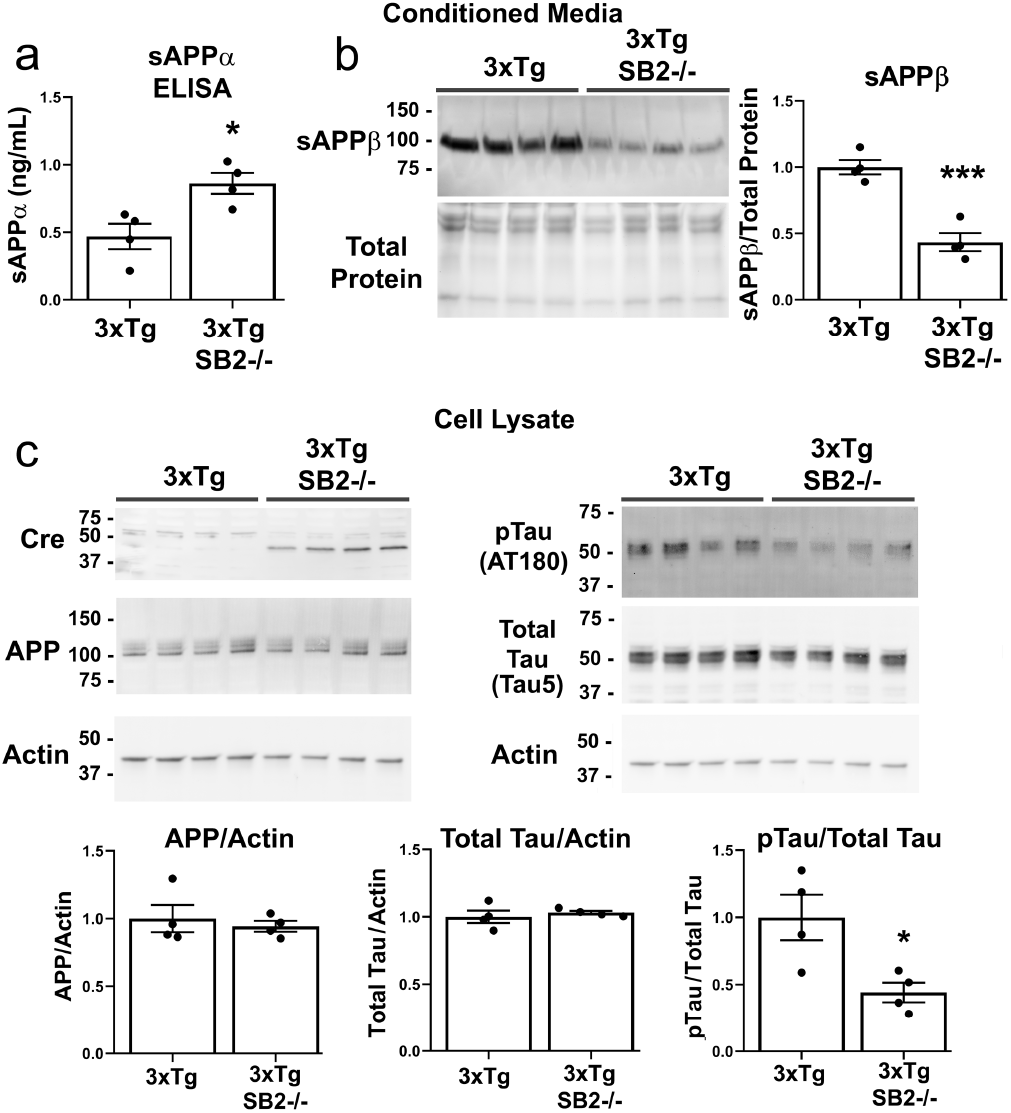
Astrocyte cholesterol regulates APP and Tau processing, not protein expression. Primary embryonic cultures of mixed neurons and astrocytes were grown by crossing 3xTg SREBP2^flox/flox^ mice with 3xTg SREBP2^flox/flox^ GFAP-Cre mice. Each culture was grown independently from a single embryo and Cre genotype was confirmed by PCR. (a) Human sAPPα levels (from transgenic APP) were measured in conditioned media from mixed cultures. (b) Protein in conditioned media was precipitated and sAPPβ, an APP fragment produced by β-secretase cleavage, was measured by western blot. (c) Mixed cultures were lysed and protein content was examined by western blot. Data are expressed as mean ± s.e.m n=4 per genotype. *P <0.05, ***P<0.001

### Activation of astrocyte cholesterol synthesis by inflammatory cytokines

The specific role of astrocytes on APP localization and Aβ plaque formation led us to consider additional signals in disease states that might work through astrocytes to upregulate their formation. Among likely regulators, inflammation is a prominent risk factor associated with AD^52^. Neuroinflammation is characterized by the release of pro-inflammatory cytokines including interleukin 1β (IL1β), interleukin 6 (IL6) and tumor necrosis factor α (TNFα). These same cytokines are known to increase cholesterol production in peripheral tissues through SREBP2 activation^53–56^, but their effects on astrocytes have not been tested.

To test the impact of inflammation on astrocyte cholesterol synthesis, we treated primary cultured astrocytes with IL1β, IL6 and TNFα. Primary astrocyte cultures were incubated for 1 hour with 1-100 ng/ml of cytokines, then assayed for total cellular cholesterol with a fluorescent cholesterol assay (see methods). All three cytokines dose dependently increased total levels of astrocyte cholesterol (up to 120%) (Figure 6a-c).

**Figure 6.**
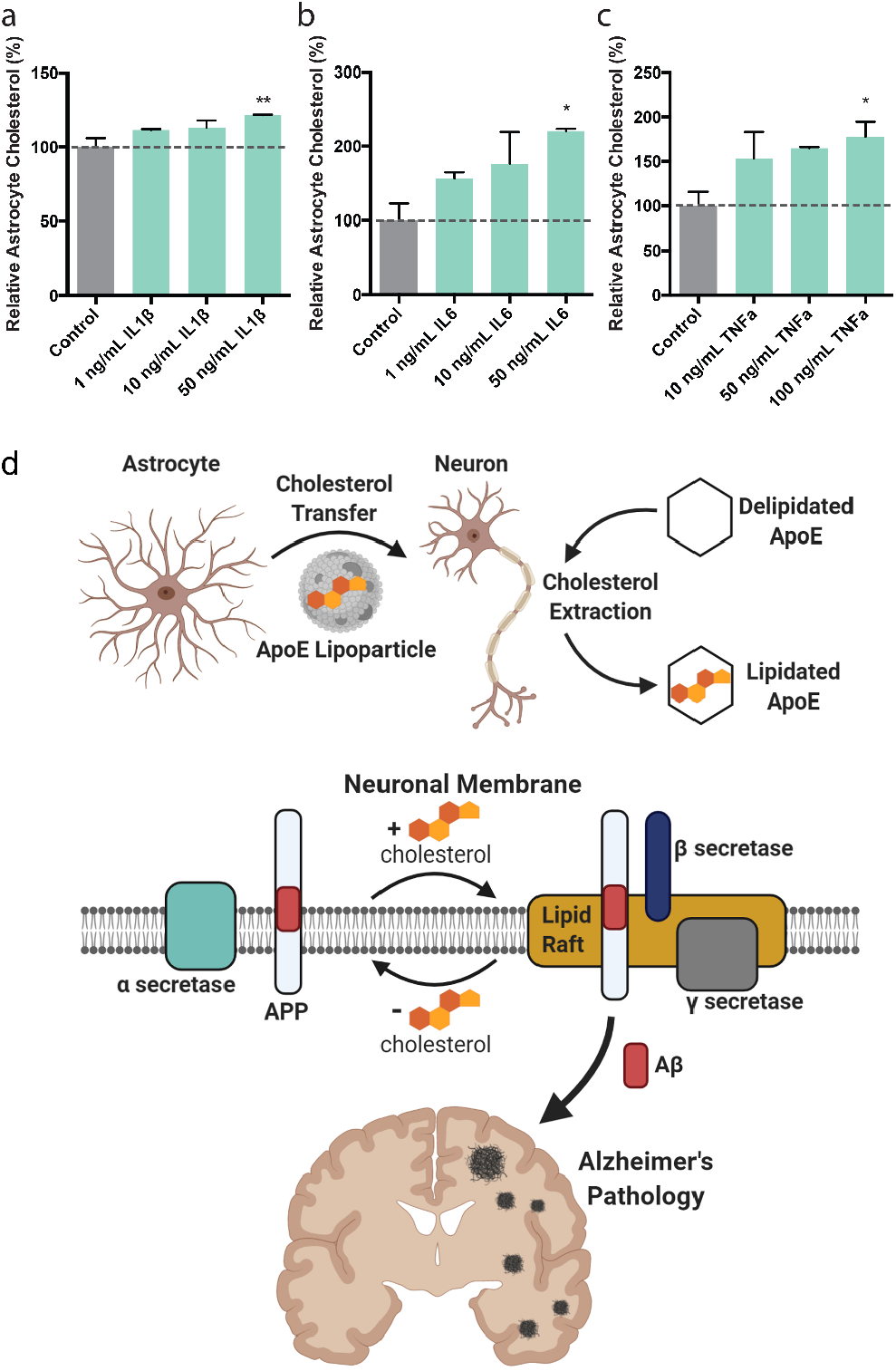
Pro-inflammatory cytokines induce cholesterol synthesis in astrocytes. (a) Interleukin-1β (IL-1β), (b) Interleukin-6 (IL-6), and (c) Tumor necrosis factor-alpha (TNFa) induce cholesterol synthesis in primary cultured astrocytes, dose dependently measured by a fluorescent based cholesterol assay. Data are expressed as mean ± s.e.m., *P<0.05, **P<0.01, one-way ANOVA. (d) A model for amyloid production in the Alzheimer’s brain. Cholesterol is synthesized in astrocytes and shuttled to neurons in ApoE lipoprotein particles. Adding recombinant cholesterol-containing apoE enriches neuronal membrane cholesterol levels, while delipidated ApoE reduces membrane cholesterol. Cholesterol loading of the neuronal membranes regulates Aβ production by increasing APP interactions with β and γ secretase. In low cholesterol membranes APP interacts with α-secretase, generating sAPPα. When neuronal membranes are loaded with cholesterol, APP increasingly interacts with β- and γ-secretases generating Aβ peptides resulting in brain plaque formation over time.

Together, our data support astrocyte cholesterol as a key regulator of neuronal Aβ formation (Figure 6d). Inflammatory cytokines, possibly from microglia, activate astrocyte cholesterol synthesis. Astrocyte-secreted ApoE then loads cholesterol into neurons. Increased cholesterol in neurons drives APP to associate with β- and γ-secretases in lipid rafts, the association of APP with lipid rafts regulates the amount of Aβ formation, and Aβ levels dictate insoluble plaques.

## DISCUSSION

While it has been known for some time that astrocytes play an important role in brain cholesterol production and express the AD genetic risk factor apoE, the role of astrocytes in AD pathogenesis remains poorly understood^57–59^. Astrocytes undergo robust morphological changes in neurodegenerative models and recent research demonstrates that astrocytes undergo broad transcriptional changes early in the AD process^60^. However, whether astrocytes are simply reacting to the AD neurodegenerative cascade or playing a role in promoting disease remains unclear. Here we demonstrate a direct role for astrocytes in promoting the AD phenotype through the production and distribution of cholesterol to neurons. Combined, our data establishes a molecular pathway that connects astrocyte cholesterol synthesis with apoE lipid trafficking and amyloidogenic processing of APP. The pathway establishes cholesterol as a critical lipid that controls the signaling state of a neuron. Cholesterol appears to be kept low as a mechanism to allow astrocytes to control APP presentation to proteolytic enzymes in the neuron. By keeping cholesterol low, the astrocyte can move the neuron through a concentration gradient that profoundly affects APP processing and eventual plaque formation. In essence cholesterol is set up to be used as a signaling lipid. Rather than targeting a receptor, it targets GM1 domains and sets the threshold for APP processing by altering GM1 domain function. This concept is likely important to other biological systems^61^ given the profound effect of cholesterol on human health.

The rise and fall of Aβ peptide with cholesterol is striking (Figure 4a-b) and the data support the proposed molecular mechanism in which amyloidogenic processing of APP is promoted by cholesterol increasing APP interactions with β and γ secretases (Figure S1). This finding is in agreement with prior research implicating increased brain cholesterol content as a factor which promotes AD associated amyloid pathology. Administration of statins to guinea pigs significantly reduces Aβ levels in the CSF^62^. APP/PS1 AD mice overexpressing a truncated active form of SREBP2 display an accelerated Aβ burden with aging^63^. Our findings contribute to literature demonstrating that amyloidogenic processing of APP in cultured cells occurs primarily in cholesterol rich membrane domains^14,64^ and is consistent with cholesterol unloading decreasing Aβ formation in cell culture^65^ and on the plasma membrane^66^. Cholesterol loading into cells regulates more than just amyloid, it also appears to play an essential role in the regulation of tau^50^. We observed a striking reduction in levels of phosphorylated tau in astrocyte SREBP2 knockout AD mice, which could result from reduced Aβ production or directly from cholesterol regulation. Our work is focused on cleavage of APP in neurons and primarily on the plasma membrane. Other mechanisms resulting from changes in brain cholesterol metabolism likely contribute to both amyloid production and clearance and require further investigation. Notwithstanding, astrocyte cholesterol is clearly a necessary factor for plaque formation even when APP is overexpressed.

The regulation of astrocyte cholesterol by cytokines is consistent with an antimicrobial function for Aβ (Figure 6d). The robust upregulation of Aβ in response to cholesterol suggests a similar mechanism likely occurs in peripheral tissue^67^ or with microhemorrhaging in the brain.^68^

At the molecular level, cholesterol loading regulates two functions: 1) the localization of APP to GM1 domains and 2) the total number of GM1 domains (Figure 1c-d). Palmitoylation drives raft association of the majority of integral raft proteins^7^, but β, and γ-secretases are also palmitoylated and remain localized to GM1 lipids suggesting a difference in relative affinity or additional factors that facilitate its release from GM1 domains. PIP2 is a polyunsaturated lipid that resides in the disordered region and pulls PLD2 away from GM1 lipids^15^. If PIP2 binds APP, but not the secretases, this would help explain APP’s selective trafficking away from GM1 lipids in low cholesterol. APP’s localization to PIP2 is not known, but PIP2 is an important factor in AD^69,70^.

Many channels are palmitoylated^71^ and may respond to apoE and GM1 lipid domain function similar to APP. Investigating the effect of astrocyte cholesterol on channels will be important for studying AD since many of the palmitoylated channels have profound effects on neuronal excitability and learning and memory^71^. In separate studies we saw cholesterol loading with apoE caused the potassium channel TREK-1 to traffic to lipid rafts similar to APP (Nayebosadri unpublished data)^72^ an effect that was reversed by mechanical force^73^.

Our data suggest cultured cells lacking apoE supplementation likely underestimate physiological GM1 domains size, especially those derived from the brain where cholesterol is high^74^. Not surprisingly, rafts appeared to be absent in cultured cells^75^. Not until cholesterol, supplied by astrocytes or cholesterol-containing apoE, was added were nanoscale rafts present (~100 nm). We conclude from our data, that the conditions with cholesterol are the more physiologically relevant conditions for cultured cells.

All astrocyte-targeting Cre lines also delete to some degree in neural progenitor cells. While our model also suffers from this short-coming, we have previously demonstrated that overall cholesterol synthesis by neurons increases to compensate for loss of astrocyte cholesterol^46^. Thus, the dramatic decrease in Aβ and pTau that we observe in our AD x SB2(-/-) mice cannot readily be explained by the minority of knocked-out neurons. In addition, microglia which are also able to produce apoE^47^, and are not impacted by the Cre, are unable to compensate for a lack of astrocyte cholesterol synthesis. Our data emphasize that the small changes in Aβ production we see in our cell culture models have a meaningful impact *in vivo*, where the cumulative impact of altering cholesterol delivery to neurons can be observed.

We conclude that the availability of astrocyte cholesterol regulates Aβ production by substrate presentation. This contributes to the understanding of AD and provides a potential explanation for the role of cholesterol associated genes as risk factors for AD.

## Methods

### Animals

Housing, animal care and experimental procedures were consistent with the Guide for the Care and Use of Laboratory Animals and approved by the Institutional Animal Care and Use Committee of the Scripps Research Institute and the University of Virginia. 3xTg-AD mice and B6129SF2/J controls were purchased from Jackson labs. The 3xTg-AD mice were maintained homozygous for all transgenes. SREBP2^flox/flox^ mice (a generous gift of Dr. Jay Horton at UT Southwestern, now available through Jackson labs) were crossed to the hGFAP-Cre as previously described^46^. These mice were in turn bred to the 3xTg-AD line and crossed back to homozygosity for both the 3xTg-AD transgenes and the SREBP2-flox. 3xTg-AD x SREBP2^flox/flox^ (AD x SB2flox) and 3xTg-AD x SREBP2^flox/flox^ x GFAP-Cre (AD x SB2^-/-^) are littermates. Only female mice were used for the AD crosses as the male 3xTg-AD mice have a much milder phenotype.

Aldh1l1-Cre^ERT2^ mice were purchased from Jackson labs. These mice were crossed to the SREBP2^flox/flox^ mice and the floxed allele was kept homozygous. Both male and female embryos were used from this mouse line.

### Cell culture

Neuroblastoma 2a (N2a) cells and human embryonic kidney 293T (HEK293T) cells were grown in Dulbecco’s Modified Eagle Medium (DMEM) containing 10% fetal bovine serum (FBS) and 1% penicillin/streptomycin. The cells were changed to a serum-free media 24 hours prior to experimentation unless otherwise noted.

Neurons were isolated from the cortices of embryonic day 18-21 CD1 mice. Cells were dissociated by papain digestion and plated on poly-D-lysine-coated (0.01 mg/mL) 8-well Lab-Tek II chambered cover glass (Thermo Fisher Scientific, #Z734853). Specifically, dissociated cells were plated in Minimum Essential Medium (MEM) supplemented with 5% FBS and 0.6% glucose and grown in Neurobasal medium (Thermo Fisher Scientific) supplemented with 20%B27 (Invitrogen), 1% Glutamax and 1% penicillin/streptomycin at 37 °C in 5% CO2. Cells were cultured *in vitro* for two days prior to experiment. Astrocytes were cultured in DMEM containing 10% FBS and 1% penicillin/streptomycin as described previously^76^.

#### Inducible Astrocyte SREBP2 Knockout Cultures

Astrocyte SREBP2 knockout cells with wild type neuron cocultures were grown by breeding ALDH1L1-Cre^ERT2^xSREBP2^flox/flox^ mice with SREBP2^flox/flox^ mice. Embryonic day 17 brain cortices were harvested from embryos and cultured individually, while tails from each individual embryo were collected for determination of Cre genotype. Meninges-free embryo cortices were digested in papain supplemented with DNAse for 30 minutes then digestion was terminated by 4:1 addition of sterile HBSS + 20% fetal bovine serum. Following enzymatic dissociation, cells were centrifuged at 300 x g for 3 minutes. The resulting cell pellet was gently resuspended in plating media (MEM + 5% FBS + 0.6% glucose) using a 1mL pipette. Cells were plated on 0.01 mg/mL poly-d-lysine overnight coated Nunc™ Lab-Tek™ II 8-chambered coverglass or glass coverslips in plating media and placed in a 37-degree 5% CO2 cell culture incubator. After two hours, plating media was gently removed from the cells and replaced with neurobasal media supplemented with 20% B27 and 1% glutamax. After 4 days in culture, astrocyte SREBP2 ablation was induced by addition of 100 nM 4-OH tamoxifen (Sigma-Aldrich H7904) to culture media for 48 hours. After 48 hours 4-OH tamoxifen was removed and media was replaced with fresh neurobasal media containing 20% B27 and 1% glutamax. Cells were then maintained in culture for 4 days and fixed for imaging as described below.

#### 3xTg Astrocyte SREBP2 Knockout Cultures

Mixed cultures of 3xTg neurons with WT or SREBP2 knockout astrocytes were generated from embryos by breeding 3xTg SREBP2^flox/flox^ mice with 3xTg SREBP2^flox/flox^ hGFAP-Cre mice. On embryonic day 17, the embryos were harvested and the cortices and tail of each embryo were collected for cell culture and PCR genotyping respectively. The mixed cultures were grown from individual embryonic cortices as performed in the inducible system described above, however cells were seeded at a high density (approximately 200,000 cells/well) to allow for a high density of neurons and astrocytes. After 3 days in culture, the cell culture media was exchanged for fresh supplemented neurobasal media. The culture media and cell lysate were then collected for ELISA and western blot experiments. Conditioned media was concentrated for western blot by addition of 250 μL trichloroacetic acid (1.42g/mL H2O) to 1 mL of conditioned media. Protein was precipitated for 20 minutes on ice and pelleted by centrifugation at 20,000 RCF at 4 degrees C. The pellet was then washed twice with ice cold acetone and resuspended in Laemmli buffer under reducing conditions.

### dSTORM Super-resolution imaging

#### Fixed cell preparation

N2a cells were grown to 60% confluence and then allowed to differentiate overnight in serum free media. Primary cortical cells were grown for two days *in vitro*. Cells were incubated with 4 μg/mL purified apoE protein for one hour in media with or without FBS supplementation. Cells were rinsed with PBS and then fixed with 3% paraformaldehyde and 0.1% glutaraldehyde for 15 min to fix both proteins and lipids. Fixative chemicals were reduced by incubating with 0.1% NaBH4 for 7 min with shaking followed by three times 10 min washes with PBS. Cells were permeabilized with 0.2% Triton X-100 for 15 min and then blocked with a standard blocking buffer (10% bovine serum albumin (BSA) / 0.05% Triton in PBS) for 90 min at room temperature. For labelling, cells were incubated with primary antibody for 60 min in 5% BSA / 0.05% Triton / PBS at room temperature followed by 5 washes with 1% BSA / 0.05% Triton / PBS for 15 min each. Secondary antibody was added in the same buffer as primary for 30 min at room temperature followed by 5 washes as stated above. Cells were then washed with PBS for 5 min. Cell labelling and washing steps were performed while shaking. Labeled cells were then post-fixed with fixing solution, as above, for 10 min without shaking followed by three 5 min washes with PBS and two 3 min washes with deionized distilled water.

#### Brain slice preparation

Mouse brain slicing and staining were performed as previously described^77^ with minor modifications. Mouse brains were fixed in 4% paraformaldehyde, incubated in a 20% sucrose/PBS solution at 4 °C for 3 days, and embedded in Tissue-Tek OCT compound (Sakura). Sagittal sections (50 μm) were collected and placed into 24-well plate wells containing PBS. Fixative chemicals were reduced by incubating with 0.1% NaBH4 for 30 min while gently shaking at room temperature followed by three times 10 min washes with PBS. Samples were permeabilized with 0.2% Triton X-100 for 2 hours and then blocked with a standard blocking buffer (10% bovine serum albumin (BSA) / 0.05% Triton in PBS) for 6 hours at room temperature. For labelling, samples were incubated with primary antibody for 3 hours in 5% BSA / 0.05% Triton / PBS at room temperature then 3 days at 4 °C followed by 5 washes with 1% BSA / 0.05% Triton / PBS for 1 hour each. Secondary antibody was added in the same buffer as primary for 3 days at 4 °C followed by 5 washes as stated above. Sample labelling and washing steps were performed while shaking. Labeled brain tissues were then post-fixed with fixing solution, as above, for 1 hour without shaking followed by three 30 min washes with PBS and two 30 min washes with deionized distilled water. Brain slices were mounted onto the 35 mm glass bottom chamber (ibidi, #81158) and 2% agarose were pipetted onto the slice to form a permeable agarose pad and prevent sample movement during imaging.

#### dSTORM imaging

Images were recorded with a Zeiss ELYRA PS.1 microscope using TIRF mode equipped with a pil-immersion 63x objective. Andor iXon 897 EMCCD camera was used along with the Zen 10D software for image acquisition and processing. The TIRF mode in the dSTORM imaging provided low background high-resolution images of the membrane. A total of 10,000 frames with an exposure time of 18 ms were collected for each acquisition. Excitation of the Alexa Fluor 647 dye was achieved using 642 nm lasers and Cy3B was achieved using 561 nm lasers. Cells and brain tissues were imaged in a photo-switching buffer comprised of 1% β-mercaptoethanol (Sigma, #63689), oxygen scavengers (glucose oxidase (Sigma, #G2133) and catalase (Sigma, #C40)) in 50mM Tris (Affymetrix, #22638100) + 10mM NaCl (Sigma, #S7653) + 10% glucose (Sigma, #G8270) at pH 8.0. Sample drift during acquisition was corrected by an autocorrelative algorithm.

Images were constructed using the default modules in the Zen software. Each detected event was fitted to a 2D Gaussian distribution to determine the center of each point spread function plus the localization precision. The Zen software also has many rendering options including removing localization errors and outliers based on brightness and size of fluorescent signals. Pair correlation and cluster analysis was performed using the Statistical Analysis package in the Vutara SRX software. Pair Correlation analysis is a statistical method used to determine the strength of correlation between two objects by counting the number of points of probe 2 within a certain donut-radius of each point of probe 1. This allows for localization to be determined without overlapping pixels as done in traditional diffraction-limited microscopy. Raft size estimation was calculated through cluster analysis by measuring the area of the clusters comprising of more than 10 particles with a maximum particle distance of 0.1 μm.

### Fluorescence activated cell sorting (FACS) of live cells

Live cell staining for FACS was performed as previously described^78^. Immediately after dissection and dissociation, cortical cells from embryonic mice were labelled with Brilliant Violet 421™ anti-mouse CD90.2 (Thy1.2) antibody (BioLegend, #140327). Cells were incubated with antibody at 4 °C for 20 minutes followed by washing with wash buffer (Ca^2+^/Mg^2+^ free PBS, 1mM EDTA, 25mM HEPES, 5% FBS, 10 units/mL DNase II). Cells were filtered through a 40-μm nylon filter (Corning, #431750) and sorted in a BD Biosciences FACSAria3 sorter. Thy1.2+ neurons were collected and cultured as described above.

### Live raft affinity assay

Modulation of raft integrity was detected by a biochemical assay based on the activity of a palmitate mediated raft localized enzyme phospholipase D (PLD), as described previously^17^. Briefly, N2a cells were seeded into 96-well flat culture plates with transparent-bottom (Corning™ Costar™, #3585) to reach confluency (~ 5 x 10^4^ per well). Then the confluent cells were differentiated with serum-free DMEM for a day and washed with 200 μL of PBS. The raft integrity assay reactions were promptly begun by adding 100 μL of working solution with or without apoE, (BioVision, #4696). The working solution contained 50 μM Amplex red, 1 U/mL horseradish peroxidase, 0.1 U/mL choline oxidase, and 30 μM dioctanoyl phosphatidylcholine (C8-PC). ApoE was directly dissolved into the working buffer from freshly made stocks before assay reagents were added. The PLD activity and the background (lacking cells) was determined in triplicate for each sample by measuring fluorescence activity with a fluorescence microplate reader (Tecan Infinite 200 PRO, reading from bottom) for 2 hours at 37°C with excitation wavelength of 530 nm and an emission wavelength of 585 nm. Subsequently, PLD activity was normalized by subtracting the background and to the control activity. Data were then graphed (Mean ± s.e.m.) and statistically analyzed (oneway ANOVA) with GraphPad Prism software (v6.0f).

### Cholesterol assay

To measure the relative changes in cellular cholesterol level after apoE or cytokine treatment, we developed an Amplex Red-based cholesterol assay modified from the raft integrity assay described above. Briefly, N2a cells or astrocytes were seeded into 96-well flat culture plates with transparent-bottom to reach confluency (~ 5 x 10^4^ per well). Then the confluent cells were differentiated with serum-free DMEM for a day. Cells were incubated with fresh DMEM for 1 hour followed by 1 hour of incubation in 100 μL of DMEM medium with or without treatment. Cytokines used in this study include: IL1β (Sigma-Aldrich, #SRP8033) and IL6 (Fisher, #PHC0064). After washing with 200 μL of PBS, cholesterol assay reactions were promptly begun by adding 100 μL of working solution containing 50 μM Amplex red, 1 U/mL horseradish peroxidase, 2 U/mL cholesterol oxidase and 2 U/mL cholesterol esterase in PBS. Relative cholesterol concentration and the background (lacking cells) was determined in triplicate for each sample by measuring fluorescence activity with a fluorescence microplate reader (Tecan Infinite 200 PRO, reading from bottom) for 2 hours at 37°C with excitation wavelength of 530 nm and an emission wavelength of 585 nm. Subsequently, cholesterol level was normalized by subtracting the background and to the control activity. End point cholesterol signals were then graphed (Mean ± s.e.m.) and statistically analyzed (one-way ANOVA) with GraphPad Prism software (v6.0f).

### ELISA

To measure the relative changes in β- and α-cleavage products of APP, N2a cells were incubated with apoE with and without FBS supplementation for 1 hour, then washed with PBS once and incubated with PBS for 1 hour. Supernatants were harvested and analyzed with Aβ40 and sAPPα ELISAs.

Relative quantitation of secreted Aβ40 was performed with commercial human Aβ40 ELISA kit (Invitrogen, #KHB3481) following the manufacturer’s instructions. For sAPPα ELISA, 5 μg/mL rabbit anti-APP antibody (abcam, #ab15272) was coated on immunoassay plates (Corning™ Costar™ Cell Culture 96 well plates, #3585) at 4 °C overnight. Then, the plates were washed with PBS three times. Next, 50 μL of supernatant was added and incubated at room temperature for 1 hour. After this, 50 μL of 2 μg/mL mouse anti-human sAPPα IgG monoclonal antibody (IBL, #11088) was added as primary detection antibody. After incubation for 3 hours at room temperature, the plates were washed with PBST buffer (PBS with 0.05% Tween-20) and then 100 μL of 100 ng/mL HRP-linked rabbit anti-mouse IgG secondary antibody (Invitrogen, #31450) was added and incubated for 1 hour. An HRP substrate, chromogen (Invitrogen, #KHB3481) was added and incubated at room temperature in the dark for 30 minutes. The substrate development was terminated by adding 100 μL of stop solution from ELISA kit (Invitrogen, #KHB3481). Relative sAPPα concentration was determined by measuring absorbance at 450 nm using a microplate reader (Tecan Infinite 200 PRO).

#### Amyloid and p-Tau ELISA from Mouse Hippocampus

Human amyloid beta 1-40 (R&D Systems DAB140B) and 1-42 (R&D Systems DAB142) was quantified in mouse hippocampus tissue using commercially available ELISA kits per manufacturer’s instructions. 60 week old female mouse whole hippocampus tissue was homogenized in RIPA buffer (Bioworld, 42020024-2) with protease inhibitors (Thermo Fisher Scientific, 78430) using a Bullet Blender® (Next-Advance) with zirconium oxide beads. The homogenate was centrifuged at 15,000 x g for 10 minutes, and the supernatant was collected to yield RIPA soluble amyloid. The remaining tissue pellet was then resuspended in 5 M guanidine-HCl diluted in 50 mM Tris, pH 8.0 with protease inhibitor and mechanically agitated at room temperature for 4 hours to extract the RIPA insoluble amyloid. Guanidine samples were diluted 1:10 in sterile PBS and centrifuged at 16,000 x G for 20 minutes. Supernatant was then collected to yield insoluble amyloid. Both soluble and insoluble amyloid supernatants were then assessed for total protein content using the Pierce 660 protein assay (Thermo Fisher Scientific, 22660). All samples were then diluted to equal concentrations of total protein. Prior to ELISA, samples were diluted 1:10 to prevent oversaturation of ELISA signal. Human phospho-tau [pT181] was measured using a commercially available ELISA per manufacturer’s instructions (Thermo Fisher Scientific, KHO0631). Guanidine extracted supernatants of hippocampus homogenates described above were utilized for the pTau ELISA.

### Confocal imaging on brain slices and primary cells

Mouse brains were fixed in 4% paraformaldehyde, incubated in a 20% sucrose/PBS solution at 4 °C for 3 days, and embedded in Tissue-Tek OCT compound (Sakura). Sagittal sections (50 μm) were collected on Superfrost/Plus slides and immunostained with mouse NF-M antibody (Santa Cruz, #sc-398532), Alexa Fluor-647-conjugated CTxB (Invitrogen, #C34778) and Cy3B-conjugated anti-mouse secondary antibody. Human amyloid beta was detected using a polyclonal antibody from Cell Signaling (#8243) with Alexa Fluor 594 antirabbit secondary (Thermo Fisher Scientific, #A21207). Primary cortical cells were fixed and labelled with Alexa Fluor-488-conjugated mouse GFAP antibody (Biolegend, #644704), or rabbit GFAP antibody (Millipore, #MAB360) with Alexa Fluor 594 anti-rabbit secondary (Thermo Fisher Scientific, #A21207). SREBP2 knockout was confirmed by measuring the downstream enzyme FDFT1 (Abcam, #ab195046, Alexa Fluor 488 secondary Thermo Fisher Scientific #A32766) pTau was imaged in brain sections using the mouse monoclonal AT180 antibody from Thermo Fisher Scientific (#MN1040). Samples were imaged on Leica TCS SP8 confocal microscope using a 25x water emersion objective. ImageJ was used to analyze images.

### Tissue Western Blots

40-week-old mouse hippocampus tissue was homogenized in RIPA buffer with protease inhibitors using a Bullet Blender®. The homogenate was centrifuged at 15,000 x g for 10 minutes, and the supernatant was collected. Total protein content was measure using the Pierce 660 absorbance assay. Sample protein content was diluted to equal concentrations, and Laemmli buffer (Biorad, 1610737) was added to each sample for SDS-Page. SDS-Page was performed to resolve protein by molecular weight using 10 micrograms of hippocampal lysate from each sample in 4-20% acrylamide pre-cast gels (Biorad, 4568094). Resolved protein was transferred to PVDF membrane and blocked with 5% BSA solution in PBST. Primary antibodies against target proteins were applied overnight at 4 C. Membranes were then washed and fluorescent secondary antibodies were applied for 2 hours at room temperature. Following 5 washes with TBS with 0.1% tween, membranes were imaged using the Bio Rad chemidoc MP fluorescent imaging system. Primary antibodies used in this study include: APP (Y188), Abcam, #ab32136; tau, Fisher Scientific, #AHB0042; Cre, Biolegend, #908001; Actin, Biorad, #12004163; sAPPβ BioLegend #813401.

### Statistics

All statistical calculations were performed in GraphPad Prism v6.0. For the Student’s t test, significance was calculated using a two-tailed unpaired parametric test with significance defined as *P < 0.05, **P<0.01, ***P<0.001, ****P<0.0001. For the multiple comparison test, significance was calculated using an ordinary one-way ANOVA with Dunnett’s multiple comparisons test.

## Acknowledgements

We thank Andrew S. Hansen and Damon Page for aspects of experimental design, E. Nicholas Petersen for his help and discussion on the imaging and imaging analysis, Eddie Grinman and the Puthanveettil lab for mouse cortical tissue. We thank biomarker core (Columbia University, NY) and flow cytometry core (Scripps Research, FL) for technical support. This work was supported by the National Institutes of Health via a Director’s New Innovator Award to S.B.H. (DP2NS087943), R01 to S.B.H. (R01NS112534), K08 (K08DK097293) and Owen’s Family Foundation Award to H.A.F., and T32 (T32DK764627) to J.A.K. We are grateful to the Iris and Junming Le Foundation for funds to purchase a super-resolution microscope, making this study possible.

## Author Contributions

Conceptualization, S.B.H. and H.W.; Methodology, H.W., J.A.K., H.A.F. and S.B.H.; Investigation, H.W. and J.A.K; Resources, S.B.H. and H.A.F., Writing – Original Draft, H.W. and S.B.H.; Writing – Review and Editing, H.W., H.A.F., and S.B.H.; Supervision, H.A.F. and S.B.H.; Funding Acquisition, H.A.F. and S.B.H.

## SUPPLEMENTAL FIGURES

**Figure S1.**
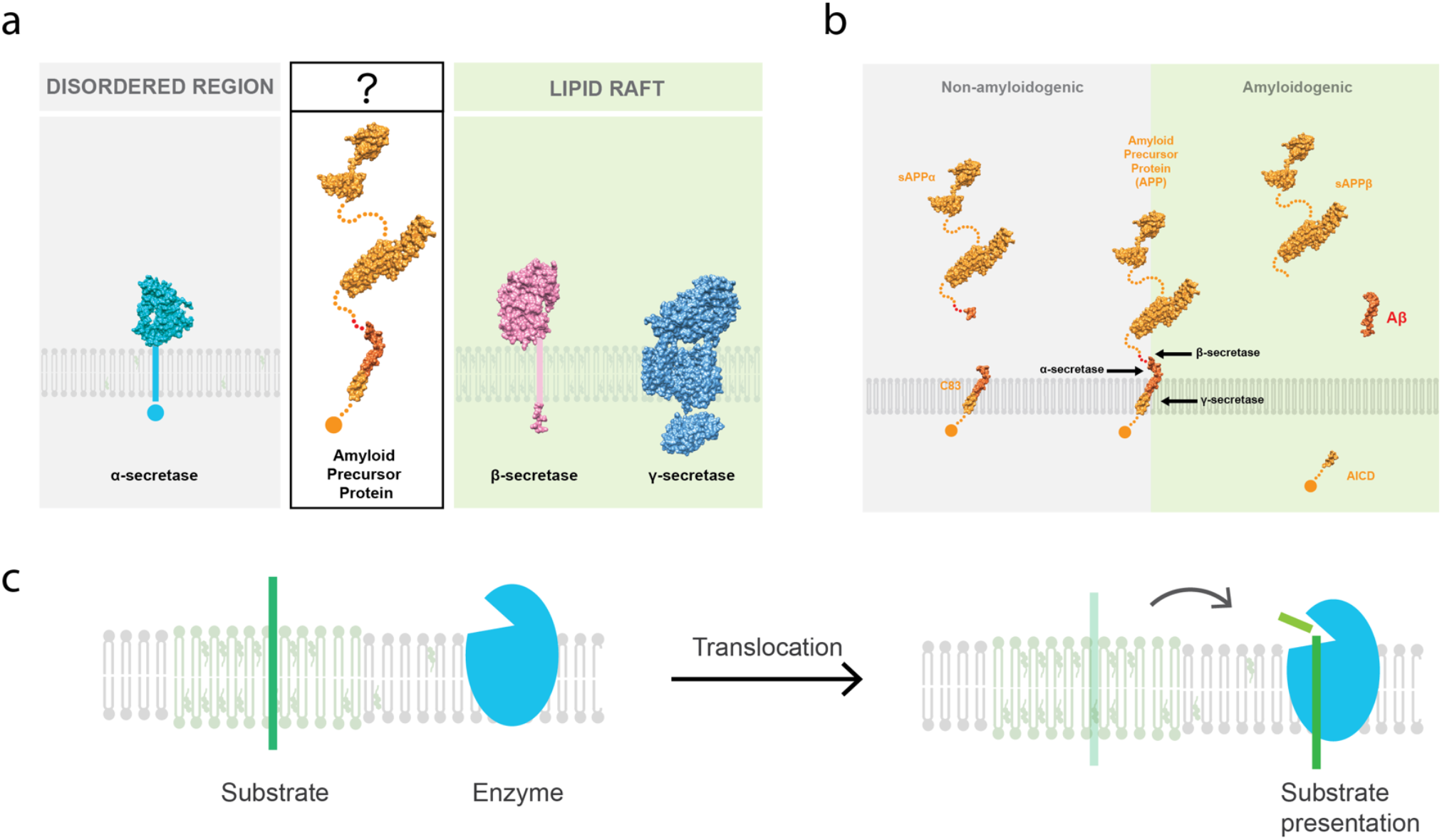
Amyloid proteins and substrate presentation. (a) The atomic structures of amyloid proteins and their predicted sub-membrane localizations based on palmitate mediated localization. Amyloid precursor protein (APP) and β- and γ-secretases are palmitoylated on their N-terminus. The palmitoylation is known to cause partitioning into detergent resistant membrane and raft-associated (green shading). α-secretase is not palmitoylated and is expected to reside in the disordered regions (grey shading). (b) The known APP cleavage sites for APP-processing proteins. In general, the α-secretase or β-secretase cleaves the APP extracellular domain generating the soluble N-terminal fragments (e.g. sAPPα or sAPPβ) and membrane-associated C-terminal fragments (CTF). CTFβ is further cleaved by the γ-secretase within the transmembrane domain releasing p3 peptide or β-amyloid (Aβ) peptide (respectively). (c) Substrate presentation is a biological process that activates a protein. The protein is sequestered away from its substrate and then activated by release and exposure of the protein to its substrate.

**Figure S2.**
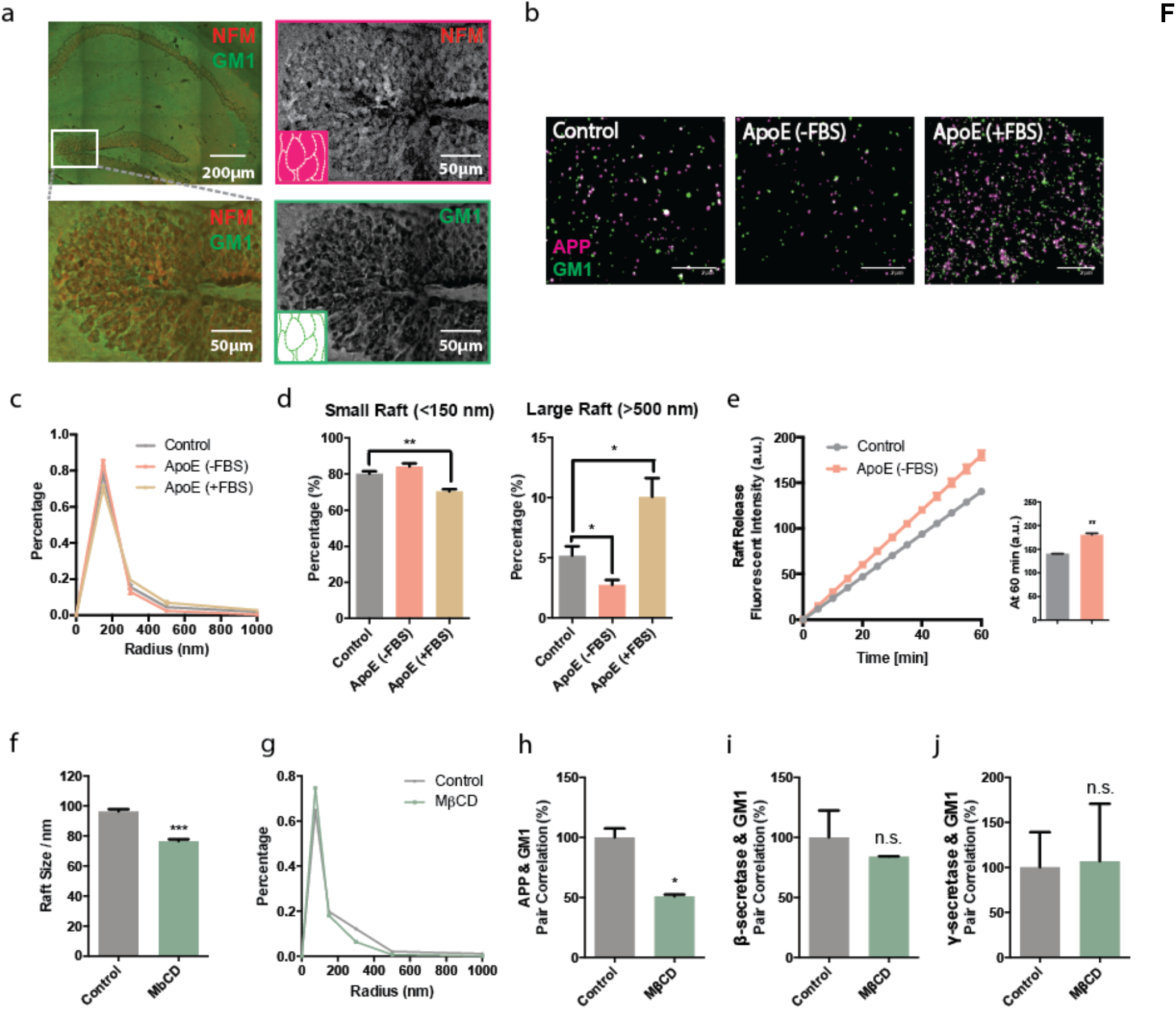
Super resolution imaging of enzymes and substrate localization with MβCD and apoE. (a) Confocal imaging on mouse brain showing GM1 lipids in plasma membrane of neurons. Neurons are labelled with neurofilament medium chains (NFM) antibody (red) and GM1 lipids are labelled with fluorophore-conjugated cholera-toxin B subunit (CTxB, green). Representative images showing NFM filled the cell bodies of hippocampal neurons while GM1 lipids are in the plasma membrane. (b) dSTORM imaging shows the effect of apoE on membrane protein’s raft association (APP in magenta, GM1 in green) in N2a cells under low (no FBS) and high cholesterol (with FBS) conditions. Scale bar is 2 μm. (cd) ApoE mediates a shift in distribution from small rafts (<150 nm) to larger rafts (>500 nm) under high cholesterol condition and opposite effect in low cholesterol condition; one-way ANOVA. (e) Raft release assay showing treatment of N2a cells with apoE causes a palmitoyl reporter enzyme to exit the raft and produce fluorescent product. Data are expressed as mean ± s.e.m., two-sided Student’s t-test *P<0.05, **P<0.01, ***P<0.001, ****P<0.0001. (f) Apparent raft size of neuroblastoma 2a (N2a) cells before and after treatment with methyl-β-cyclodextrin (MβCD). (g) Cholesterol extraction by MβCD mediates a shift in raft size distribution from large rafts (>500 nm) to smaller rafts (<150 nm). (h) Lipid raft disruption by MβCD moves APP from lipid rafts into disordered region. (i-j) β- and γ-secretase’s lipid raft localization are insensitive to MβCD-mediated raft disruption.

**Figure S3.**
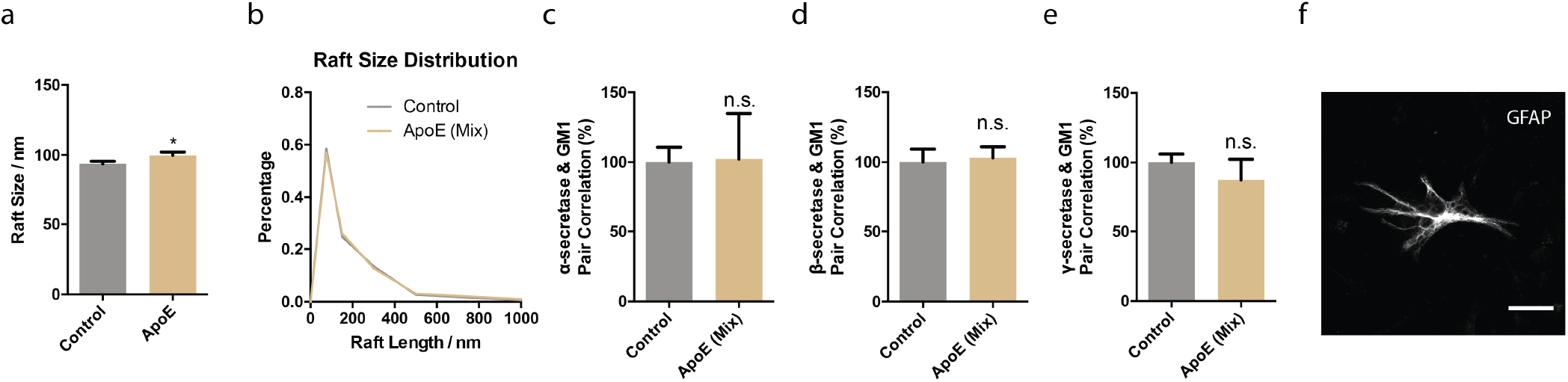
ApoE’s effect on secretases in cortical neurons. (a) Apparent raft size of primary cortical neurons before and after treatment with apoE in mixed culture with astrocytes. (b) Raft size distribution analysis showing apoE mediates a shift in distribution from small rafts (<150 nm) to larger rafts (>500 nm) in cortical neurons when culturing with astrocytes. (c-e) α-, β- and γ-secretase lipid raft localization in cortical neurons do not respond to apoE-mediated raft modulation when culturing with astrocytes. Data are expressed as mean ± s.e.m., n=3-10, two-sided Student’s t-test. (f) Astrocyte in mixed cortical culture is labelled with GFAP antibody.

**Figure S4.**
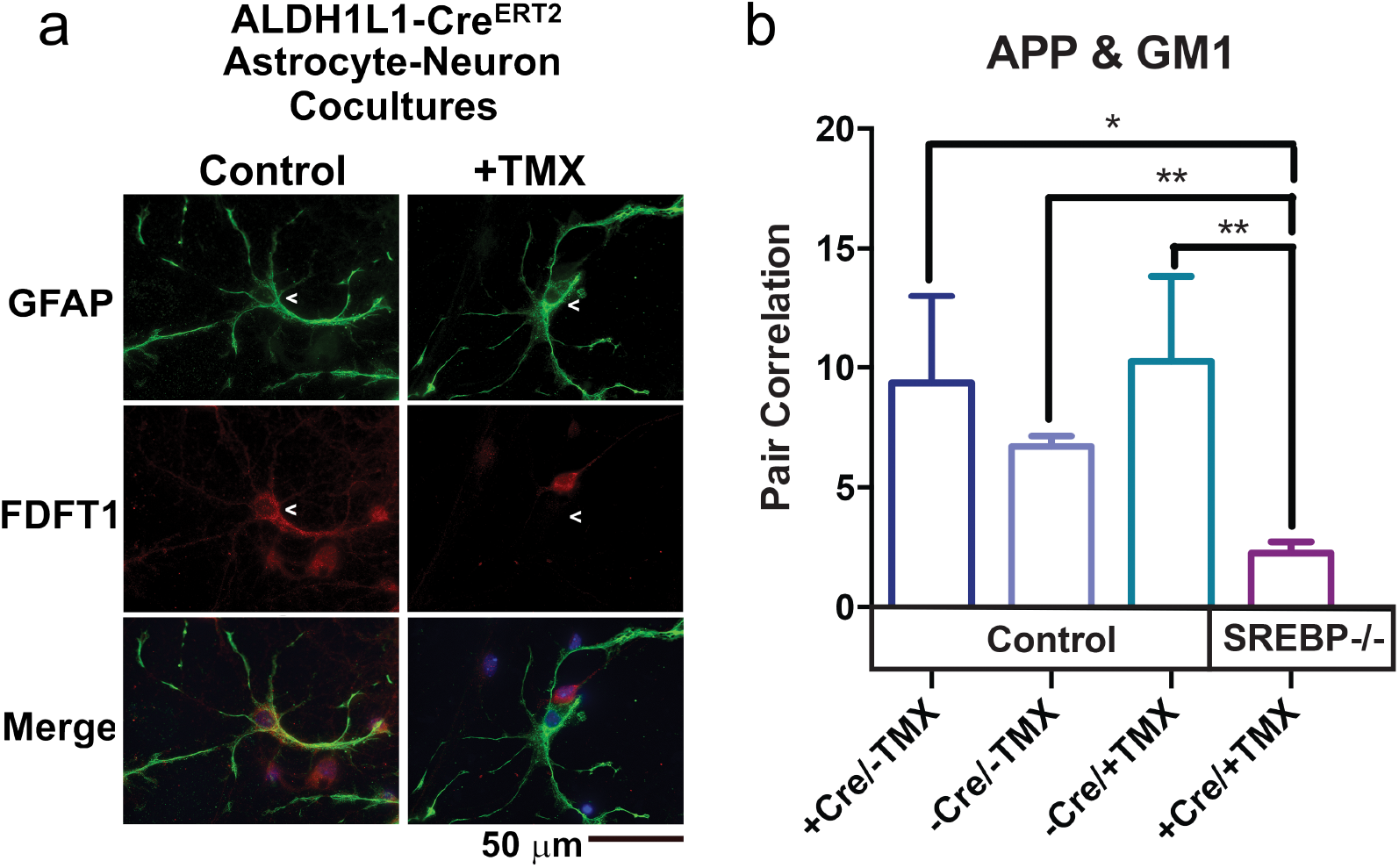
Induction of astrocyte SREBP2 gene ablation disrupts the cholesterol synthesis enzyme FDFT1 and abolishes APP and GM1 pair correlation. (a) Astrocyte-neuron co-cultures are grown from SREBP2^flox/flox^ Aldh1L1-Cre^ERT2^ embryos. Astrocyte SREBP2 knockout was induced by application of 4-hydroxytamoxifen (TMX) to cultures. Following knockout, cells were immunostained for the cholesterol synthesis enzyme FDFT1 and the astrocyte marker GFAP and imaged. (b) Mixed primary cultures of neurons and astrocytes derived from SREBP2^flox/flox^ x Aldh1L1-Cre and Cre negative littermates were treated with 4-hydroxytomoxifen (TMX) to delete SREBP2. APP lipid raft localization in cortical neurons is lost when SREBP2 is knocked out of astrocytes. TMX does not change APP localization in Cre negative cells.

**Figure S5.**
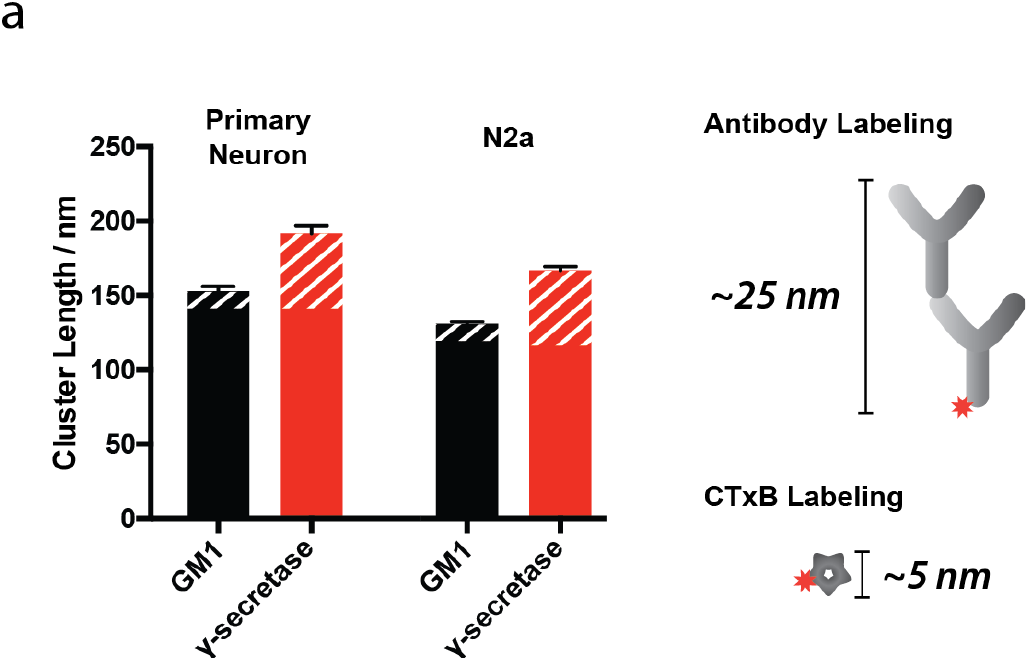
Analysis of raft size. (a) Comparison of the size of CTxB labeled GM1 domains from primary neurons and N2a cells with antibody labeled γ-secretase. The potential added diameter from antibody or CTxB labeling is indicated by white stripes. Fixed γ-secretase within the GM1 lipids form clusters approximately the same size as CTxB labeled GM1 clusters, suggesting CTxB does not cluster unfixed GM1 lipids after treatment^80^.

**Figure S6.**
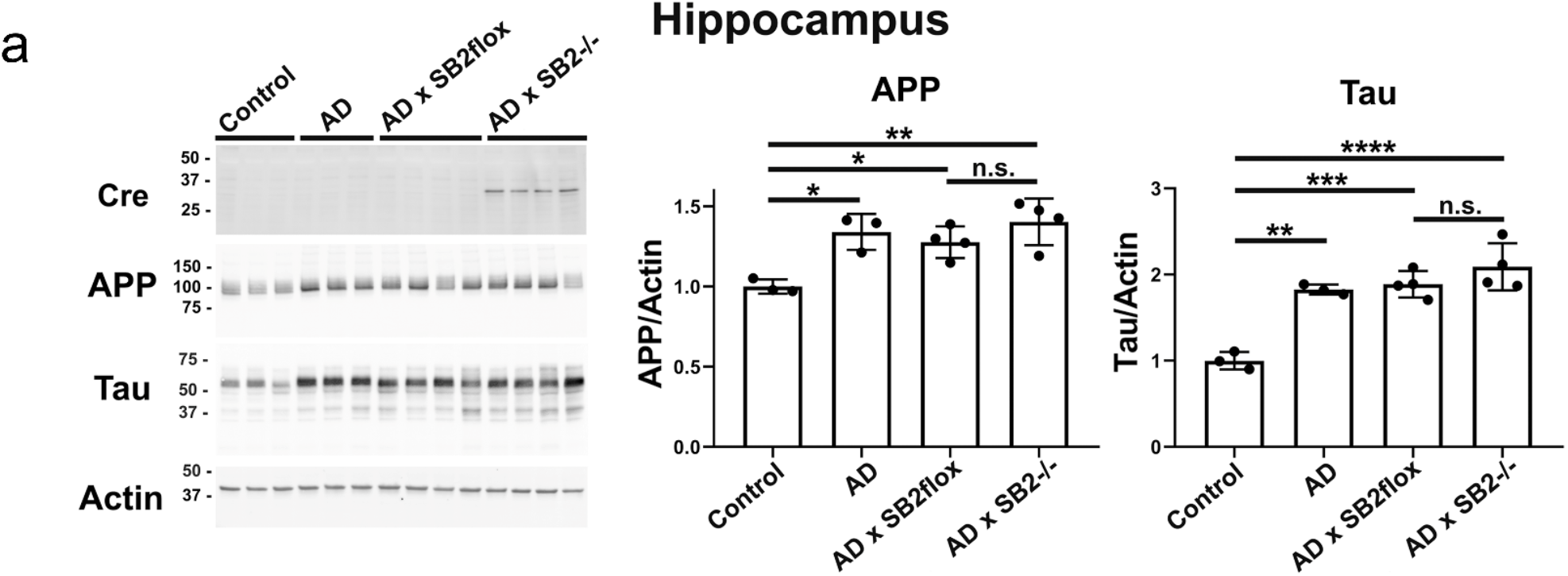
Loss of astrocyte SREBP2 in the 3xTg AD models does not impact total transgene protein levels in the hippocampus. 3xTg-AD (AD) mice express transgenes for mutant human amyloid precursor protein (APP) and tau. The AD mice were crossed to SREBP2^flox/flox^ mice (AD x SB2flox) or to SREBP2^flox/flox^GFAP-Cre^+/-^ mice (AD x SB2^-/-^). Control mice do not express the human transgenes. Animals were aged to 40 weeks and hippocampus tissue was dissected and homogenized for western blot. Data are expressed as mean ± s.e.m, n=3-4 animals per genotype. *P<0.05, **P<0.01, ***P<0.001, ****P<0.0001, one-way ANOVA with Tukey’s post hoc analysis.

